# Gene expression profiling reveals B cells are highly educated by the pancreatic environment during autoimmune diabetes

**DOI:** 10.1101/2022.05.19.492647

**Authors:** Joanne Boldison, Jessica Hopkinson, Joanne Davies, James A Pearson, Pia Leete, Sarah Richardson, Noel G Morgan, F. Susan Wong

**Author notes:** Corresponding author: J. Boldison, Institute of Biomedical & Clinical Science, University of Exeter, RILD Building (Level 4), Barrack Road, Exeter, EX2 5DW.

## Abstract

B cells play an important role in driving the development of type 1 diabetes, however, it remains unclear how they contribute to local beta-cell destruction during disease progression. Using gene expression profiling of B cell subsets in the pancreas and pancreatic lymph nodes, we reveal that B cells are highly modified by the inflamed pancreatic tissue and can be distinguished by their transcriptional profile from those in the lymph node. We identified both a discrete and a core shared gene expression profile in islet CD19^+^CD138^-^ and CD19^+^CD138^+^ B cell subsets, the latter known to have enriched autoreactivity during diabetes development. Upon localisation to pancreatic islets, CD138^+^ B cells overexpressed genes associated with adhesion molecules and growth factors compared to CD138^-^ B cells. Their shared signature displayed gene expression changes related to the differentiation of antibody-secreting cells and gene regulatory networks associated with interferon signalling pathways, pro-inflammatory cytokines and toll-like receptor activation. Finally, abundant TLR7 expression was detected in islet B cells, and was enhanced specifically in CD138^+^ B cells. Our study, therefore, provides a detailed transcriptional analysis of islet B cells identifying specific gene signatures and interaction networks that point towards a functional role for B cells in driving autoimmune diabetes.

## Introduction

Type 1 diabetes is an organ-specific autoimmune disease, characterised by immune-mediated beta-cell destruction, resulting in insulin deficiency. Current therapeutic goals include attempts to slow or halt disease progression using immune-targeted reagents and, among these, B cell depletion therapy (Rituximab) has been successful in delaying the loss of C-peptide, albei temporarily [1]. The success achieved in clinical trials when B cells are depleted or functionally compromised, is mirrored in numerous animal studies [2, 3]. Immunohistological analysis has revealed that CD20^+^ B cells are present in the pancreas of individuals with type 1 diabetes [4, 5], and that their frequency correlates with age at diagnosis. Younger children (<13y at onset) have the greatest proportion of infiltrating CD20^+^ cells and a more aggressive disease [6], highlighting the need for further interrogation of the role of B cells in the immune-mediated destruction of pancreatic beta cells.

Studies in the non-obese diabetic (NOD) mouse model demonstrate that B1 B cells are important in the initiation of type 1 diabetes [7, 8], being present in the pancreas early in the disease process, whereas established islet B cells have a more follicular phenotype [9]. During established insulitis, B cells become CD5 negative and can upregulate CD138 (Syndecan-1) [10, 11]. Recently, we have described distinct populations of islet-specific B cells in NOD mice that can be grouped according to the expression of CD138, IgD and CD19 [11, 12]. Importantly, insulin-specific autoreactive B cells have an intermediate expression level of CD138, and exist as either CD138^int^CD19^+^ or CD138^int^CD19^-^ [11]. In addition, a small subset of a highly proliferative B cells with a high expression of CD138 and CD44, and lower expression of CD19 can be distinguished [12], which resembles a murine plasmablast phenotype [13].

In this study, we have investigated the gene expression profile of distinct B cell subsets found in the pancreas in NOD mice with extensive insulitis. We carried out a comprehensive transcriptional analysis of B cells to explore their primary functional role at the local tissue level during the development of autoimmune diabetes.

## Material and Methods

### Mice

NOD/Caj mice, originally from Yale University, were bred in-house at Cardiff University. Mice were maintained at Cardiff University in specific pathogen-free isolators or scantainers. All animals received water and food *ad libitum* and were housed in a 12h dark/light cycle. The animal experiments were conducted in accordance with United Kingdom Animals (Scientific Procedures) Act, 1986 and associated guidelines. Female NOD mice aged 16-20 weeks of age were chosen at random and processed in groups for gene array experiments.

### Tissue preparation

Pancreatic lymph nodes (PLNs) were disrupted mechanically with a 30G needle. Pancreata were inflated with collagenase P solution (1.1 mg/mL) (Roche, Welwyn Garden City, U.K.) in Hanks’ balanced salt solution (with Ca2 and Mg2) through the common bile duct, followed by collagenase digestion with shaking at 37°C for 10 min. Islets were isolated by Histopaque density centrifugation (Sigma-Aldrich, Dorset, U.K.) and hand-picked under a dissecting microscope. For flow cytometric sorting, islets were then trypsinised to generate a single-cell suspension. Islet cells were rested at 37°C in 5% CO2 in Iscove’ s modified Dulbecco’s media (supplemented with 5% FBS, 2 mmol/L L-glutamine, 100 units/mL penicillin, 100 m g/mL streptomycin, and 50 mmol/L b -2-mercaptoethanol) overnight.

### Immunofluorescence

Pancreatic tissues were frozen in optimal cutting temperature (OCT) medium and sectioned at 7-10μm thickness. Pancreatic sections were fixed in 1% paraformaldehyde for 1 h at room temperature. Following fixation, tissue was permeabilized with 0.2% Triton X-100 and blocked with 5% FBS beforμe the addition of a primary antibody mix before secondary labelling was performed. Primary antibodies used were rat or rabbit anti-mouse CD20 (Abcam; Cell Signalling respectively), rat anti-mouse CD138 (Clone 281.2, BioLegend), directly conjugated goat anti-mouse IgA (Cambridge Bioscience), rabbit anti-mouse TLR7 (Novus Biologicals), biotinylated anti-insulin (clone D6C4; Abcam). Secondary antibodies used were Alexa Fluor 488 (Invitrogen), Alexa Fluor 647 (Abcam), Streptavidin-conjugated Alexa Fluor 568 (Invitrogen). All slides were mounted with VECTASHIELD mounting medium with DAPI (Vector Laboratories). All sections were imaged on a Leica DMi8 TCS SP8 Confocal 8 and processed in Fiji Image J (Version 2.0).

### Flow Cytometry and Fluorescent Activated Cell Sorting (FACS)

Cells were incubated with TruStain (anti-mouse CD16/32; BioLegend) for 10 min at 4°C followed by fluorochrome-conjugated monoclonal antibodies (mAbs) against cell surface markers for 30 min at 4°C. B cell phenotyping multiparameter flow cytometry was carried out using the following mAbs: CD19 (6D5), CD138 (281-2), B220 (RA3-6B2), IgD (11-26c.2a), IgM (RMM-1), CD3 (145-2C11), CD11c (N418), CD11b (M1/70), CD127 (A7R34). For TLR7 (A94B10, BioLegend) staining cells were fixed/permeabilized using eBioscience nuclear transcription kit. FACS was carried out with the following mAbs: CD3 (145-2C11), CD11c (N418), CD11b (M1/70), CD19 (6D5), CD138 (281-2), IgD (11-26c.2a). Dead cells were excluded from analysis by live/dead exclusion dye (Invitrogen). Cell suspensions were either acquired on an LSRFortessa (BD Biosciences) and analysed using FlowJo version 10.1 software (Tree Star, Ashland, OR) or sorted on an FACSAria III (BD Biosciences).

### RNA isolation

RNA was isolated from cells with Qiagen RNAeasy Micro kit, according to manufacturer’s instructions. B cell subsets were sorted directly into RLT buffer (Qiagen), and RNA isolated immediately. Total RNA was quantified, and quality assessed using the RNA 6000 Pico Kit on the Agilent 2100 Bioanalyzer (Agilent Technologies, Palo Alto, CA, USA).

### Clariom S array

Biotin-labelled targets for the microarray experiment were prepared using 500pg of total RNA. Amplified and biotinylated complementary RNA (cRNA) was synthesized, fragmented and labelled using the Genechip® Pico Reagent Kit (ThermoFisher Scientific) in conjunction with the Genechip® Poly-A RNA Control Kit as described in the User Manual (P/N 703210). RNA amplification was achieved using low-cycle PCR followed by linear amplification using T7 *in vitro* transcription (IVT) technology. For each sample, a hybridization cocktail of biotinylated target was incubated with a GeneChip® Mouse Clariom S array (ThermoFisher Scientific) at 60rpm for 16hours at 45°C in a Genechip® Hybridisation Oven 645 (Affymetrix). After hybridization, non-specifically bound material was removed by washing and specifically bound target was detected using the GeneChip® Hybridization, Wash and Stain Kit (ThermoFisher Scientific), in conjunction with the GeneChip® Fluidics Station 450 (Affymetrix). The arrays were scanned using a GeneChip® Scanner 3000 7G (Affymetrix) in conjunction with Affymetrix Genechip® Command Console (AGCC) software and .CEL files were generated from the resultant probe cell intensity data.

### QPCR

Total RNA was converted to cDNA using the High-Capacity RNA-to-cDNA™ kit (ThermoFisher) according to the manufacturer’s instructions. For quantitative PCR we used the TaqMan™ Fast Advanced Master Mix (ThermoFisher) according to manufacturer’s instructions. Samples were prepared using a EPMotion P5073 Liquid handling robot (Eppendorf) and amplified in duplicate alongside a housekeeping gene on a ViiA7 Real-Time PCR system (ThermoFisher). Normalisation of samples was performed by dividing the value of the gene of interest by the value of the housekeeping gene (GAPDH) (ΔCT) and an average ΔCT was presented.

### Gene array analyses

CEL files generated from Affymetrix Clariom S human arrays were imported into the Bioconductor package oligo version 1.56.0 [14] or imported into the Transcriptome Analysis Console (TAC) software (ThermoFisher). Firstly, DNA microarray analyses were performed in R version 4.0.3. Data were normalized using the Robust Multichip Average (RMA) algorithm [15]. Differential expression (DE) analysis was performed using the linear models for microarray data (limma) package version 3.48.1 [16]. Linear models were determined for each transcript cluster (gene) and an estimate for the global variance calculated by an empirical Bayes approach [17]. A moderated *t*-statistic was computed for each transcript cluster with the resulting *p*-values were corrected using the Benjamini-Hochberg (BH) method to control the False Discovery Rate (FDR). Genes were annotated using the Affymetrix ‘clariomsmousetranscriptcluster.db’ annotation data package available from Bioconductor. For further analysis the ebayes method was used in the TAC software to compare gene expression with previously published data and perform Venn analysis. The PANTHER classification system [18] or the Functional Annotation Tool DAVID [19] was used for gene ontology (GO) analysis with either upregulated or downregulated (FDR<0.05, FC >2). The statistical overrepresentation analysis tool was used in PANTHER to determine significant GO terms. Significant gene regulatory networks and pathways were investigated using Ingenuity Pathway Analysis (Qiagen Bioinformatics) and quantified with a Z-score and -log(*p*-value). All genes were uploaded and an FDR<0.05 and a fold change >1.5 threshold set. Finally, bubble plots were generated using ggplot2 Rstudio package (version 1.2.5042), hierarchical clustering heatmaps were generated in Rstudio with either ggplot2 or Pheatmap packages. Other heatmaps, volcano or scatter plots and principal component analysis (PCA) were performed in GraphPad Prism version 9.

### Statistical analysis

Statistical analyses were performed in R software or GraphPad Prism software. Genes were discarded from the analysis if differential expression failed to be significant (adjusted *p*-val <0.05, Benjamini-Hochberg correction for multiple testing). Other statistical tests used are provided in the figure legends.

## Results

### Abundance of B cell subsets in the pancreas during autoimmune diabetes development

We have described previously four islet-specific B cell subsets, differentiated according to the expression of CD138, among which a population that is CD19^+^CD138^+^ (intermediate levels of expression) is enriched in autoreactive B cells [11]. We sought to confirm and clarify the abundance of the CD19^+^CD138^-^ and CD19^+^CD138^+^ islet B cells in the pancreatic islets of NOD mice during diabetes development (Figure 1). Previously, we reported the abundance of pancreatic CD19^+^CD138^-^ (mean SEM; 7.1 1.09%) and CD19^+^CD138^+^ (mean SEM; 3.6 2.7%) B cells in younger NOD mice (6-8 weeks-old) [12]. We have now compared the frequency of pancreatic B cell subtypes in 12–18-week-old NOD mice and show a significant increase in both the CD19^+^CD138^-^ (*p*<0.01) and CD19^+^CD138^+^ (*p*<0.05) B cells, compared to young NOD mice (*p*<0.05) (Fig. 1A), and a shift towards a 1:1 ratio during the progression of islet inflammation (*p*=0.07) (Fig. 1B). At the onset of diabetes, both CD19^+^CD138^-^ and double-positive CD19^+^CD138^+^ were identified in the few remaining insulin-containing islets and in pancreatic immune cell clusters (Fig. 1C). CD138^+^ cells that did not express CD19 were also evident (Fig. 1C), corroborating our previous observations made using flow cytometry [11]. Overall, both single positive CD19^+^CD138^-^ and double-positive CD19^+^CD138^+^ B cells comprised prominent components of the immune infiltrates found in the pancreas of NOD mice with diabetes.

**Fig 1.**
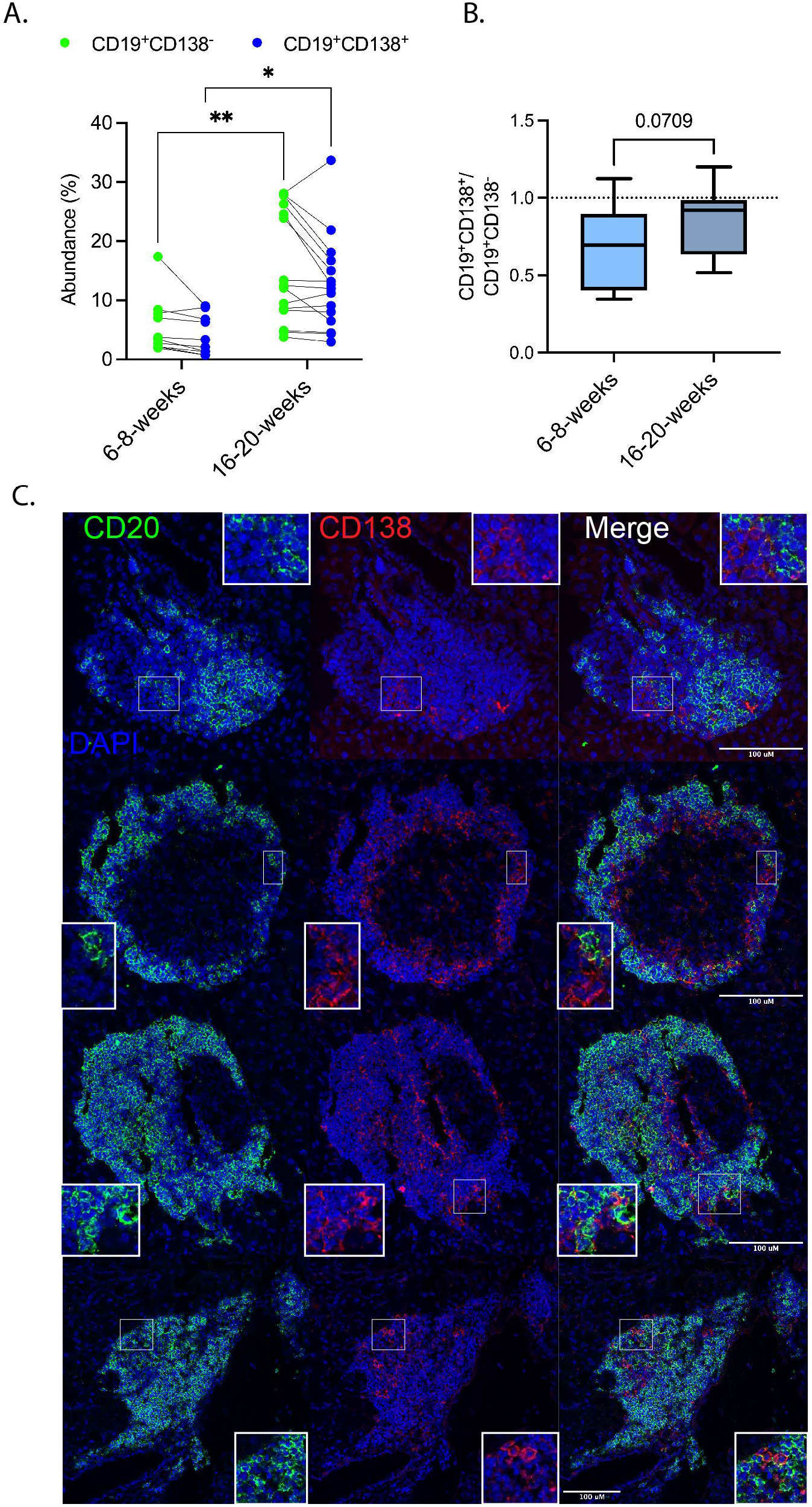
Abundance of B cell subsets in the pancreas during autoimmune diabetes development (A, B) Pancreatic islets from either single or groups of female NOD mice aged 16-20 weeks old were taken before flow cytometric analysis on CD19^+^CD138^-^ (green) and CD19^+^CD138^+^ (blue) B cells. A: Overall percentages for both CD19^+^ B cell subsets compared to younger NOD mice previously assessed [12]. Each dot represents either one mouse or pooled mice. *<0.05, **<0.01 by two-way Anova with a Bonferroni’s multiple comparison test. B: Ratio of CD19^+^CD138^+^ to CD19^+^CD138^-^ B cells. p= 0.0709 by Mann-Whitney *U* test. Data are from 5 independent experiments. CD19^+^ B cells were gated on Live singlets CD3^-^CD11b^-^CD11c^-^ cells. (C) Representative images of pancreatic sections from diabetic NOD female mice (blood glucose >13.9mmol/L) stained with CD20 (green), CD138 (red) and nuclear DAPI counterstain. Images represent pancreatic islets or immune cell clusters in 3 individual mice.

### CD19^+^CD138^-^ and CD19^+^CD138^+^ B cells have a similar transcriptional profile

Previously, in NZB/W mice, splenic CD19^+^CD138^+^ cells were described as B cells with a mature phenotype akin to an early intermediary antibody-secreting cell (ASC) stage. They respond to stimuli via both TLR4 and CD40 receptors, and the population contains a small number of Ig (immunoglobulin) secreting cells [20]. We now corroborate these conclusions and show that CD19^+^CD138^+^ B cells present in the NOD spleen display a mature follicular B cell phenotype (ESM Fig.1A), in common with most B cells located in pancreatic islets during established insulitis (IgD^+^IgM^low^) [9].

To determine the transcriptional profile of the different islet-specific B cell subsets, a gene expression array was used. B cell populations (CD19^+^CD138^-^, CD19^+^CD138^+^, CD19^-^CD138^+^) were purified by flow cytometry (ESM Fig. 1B), from pancreatic islets and the pancreatic lymph nodes (PLN) of groups of 16-20-week-old female NOD mice and used for RNA isolation. Firstly, the differentially expressed genes (DEGs) were compared between the two B cell subsets (CD19^+^CD138^-^ vs CD19^+^CD138^+^) recovered from the PLN and pancreatic islets (Figure 2). Genes were determined as differentially expressed if they changed by more than 2-fold and had a false discovery rate (FDR) <0.05. By these criteria, *Sdc1 (*the gene encoding CD138) was the only differentially expressed gene (FDR <0.05) in both the PLN and pancreatic islets (heatmap, red box, Fig. 2A). The heatmap demonstrates DEGs with a non-corrected *p*-value of <0.001. We further compared the upregulated genes in the PLN and pancreatic islets, to the gene list available via the ImmGen (Immunological Genome Project) database [21]. From this analysis, we noted that CD19^+^CD138^+^ B cells were associated with more mature subsets such as germinal centre, memory and plasmablast-like B cells (ESM Fig. 1C). An ASC signature gene, *Endou*, [22] was also upregulated in these cells, although *Sdc1* was still the most induced gene in the CD19^+^CD138^+^ B cell subset.

**Fig 2.**
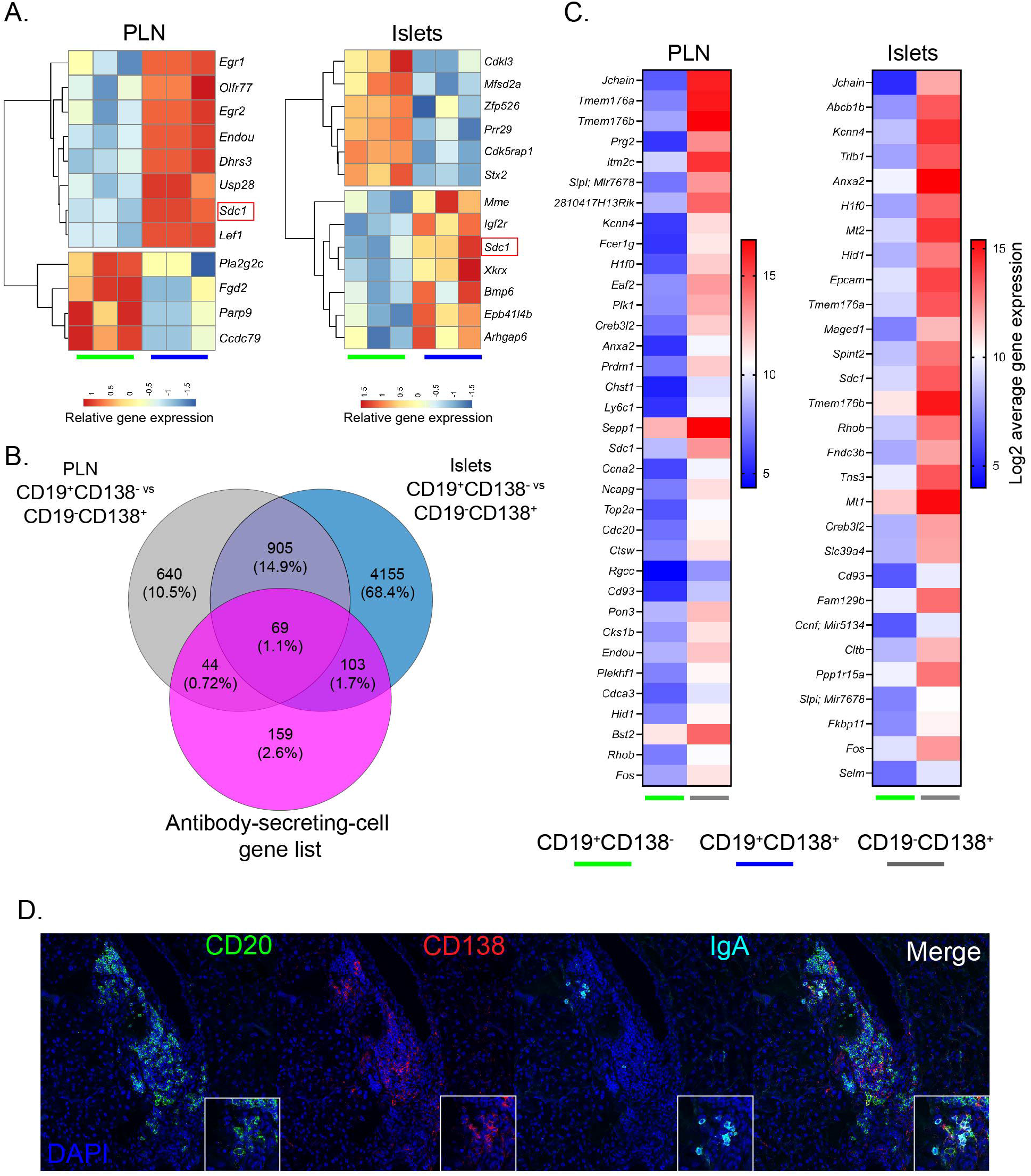
CD19^-^CD138^+^ pancreatic immune cells are enriched with fully differentiated plasma cells (A, B, C) Pancreatic lymph nodes (PLN) and pancreatic islets from groups female NOD mice aged 16–20-week-old were taken (n=8) and CD19^+^CD138^-^ (green), CD19^+^CD138^+^ (blue) and CD19^-^CD138^+^ (grey) cells were FACS sorted for RNA isolation and gene array analysis. A: Unsupervised hierarchical clustering heatmaps of the differentially expressed genes (uncorrected *p*<0.001) from the PLN (left) and pancreatic islets (right) comparing CD19^+^CD138^-^ and CD19^+^CD138^+^ B cells. Three individual sorted populations were arrayed for each B cell subset, each column represents the relative gene expression from one experimental sample. Red box highlights *Sdc1* (Sydecan1, CD138) gene expressed at FDR<0.05. B: Venn diagrams to show overlaps and differences in genes significantly (FDR<0.05, >2-fold) upregulated and downregulated between cell subsets CD19^+^CD138^-^ vs CD19^-^CD138^+^ cells in the both the PLN and pancreatic tissue, and an antibody secreting cell (ASC) gene list. C: Heatmaps showing in the PLN and pancreatic islets the top associated ASC genes differentially overexpressed in CD19^-^CD138^+^ cells compared to CD19^+^CD138^-^ sorted subsets. Average gene expression is shown from 3 individual samples. D: Representative images of pancreatic sections from diabetic NOD female mice (blood glucose >13.9mmol/L) stained with CD20 (green), CD138 (red) and IgA (cyan) and nuclear DAPI counterstain. Images represent an immune cell cluster and are representative of 3 individual mice.

### Both innate lymphocytes and fully differentiated plasma cells are enriched in the CD19^-^CD138^+^ cell subset in pancreatic islets

We have previously suggested that pancreatic immune CD138^+^ cells that do not express CD19 comprise a heterogenous mix of B cells differentiating into antibody-secreting cells (ASC) on the basis that they express CD138 and have a higher expression of CD44 and Ki67, compared to CD19^+^ B cells [12]. However, as other cell types can also express CD138 [23, 24] we sought to understand if this population were indeed enriched in mature ASC. In the PLN, CD19^+^CD138^-^ B cells were compared with CD19^-^CD138^+^ B cells in the gene array dataset (ESM Fig. 2A). A significant number of genes were differentially expressed between these populations (1658 genes, FDR *p*<0.05, >2-fold), with 1125 genes upregulated and 533 genes downregulated in the CD19^-^CD138^+^ subset (ESM Table 1). The top 20 upregulated genes (by fold change) were clustered by heatmap (ESM Fig. 2A), and this revealed that some of the highly upregulated genes (e.g. *Il7r* and *Cd7*; the former being important for early B cell maturation [25]) are abundantly expressed by innate lymphocytes [26]. Furthermore, genes such as *Klrb1b, Rora* and *Il17re* were also highly upregulated and are, again, associated with innate lymphocytes [27]. In the populations of cells recovered from pancreatic islets, these findings were recapitulated (ESM Fig. 2B, ESM Table 2).

Somewhat surprisingly, it was revealed that *Jchain* (which is considered to be expressed solely in plasma cells [20]) was highly upregulated in the CD19^-^CD138^+^ cells (compared to CD19^+^CD138^-^ cells) (ESM Fig. 2A, ESM Table 2). Other differentially expressed genes, such as *Tmem176a* and *Tmem176b*, are expressed both in plasma cells [28] and in innate lymphoid cell types [29]. Considering this, we compared the set of upregulated genes found in CD19^-^CD138^+^ cells from both PLN and pancreatic islets, with published ASC signature genes [22] and found 84 shared genes in the PLN, and 89 shared genes in the pancreatic islets (Venn diagrams, Fig. 2C). Key upregulated genes included *Prdm1* (Blimp1) and *Xbp1* (X-box binding protein 1), both of which are essential for plasma cell differentiation [30, 31]. In Figure 2D we show the top plasma cell-related genes that were changed by >7-fold (FDR<0.05) in both tissues. Since the *Jchain* gene encodes the protein component of both IgM and IgA immunoglobulins we sought to examine if IgA-secreting cells can be observed in the pancreatic tissue of NOD mice with diabetes. Occasional groups of IgA^+^ cells were seen in pancreatic immune cell clusters, with most IgA^+^ cells also expressing CD138^+^ (Fig. 2D). Thus, the CD19^-^CD138^+^ subset identified in both NOD PLN and pancreas contains cells with a more differentiated plasma-cell phenotype and a population consisting of innate-like lymphocytes.

### B cell populations are significantly modified by the inflamed pancreatic environment

Based on the discovery that CD19^+^ B cells represent the more abundant population (compared to CD19^-^CD138^+^ cells, which are not enriched as the disease progresses [12] (data not shown)), we next focused on the CD19^+^ B cells present in pancreatic tissue. To understand if B cells are altered in the inflamed pancreatic environment we compared the corresponding B cell subsets between the PLN and pancreatic islets (Figure 3). In the CD19^+^CD138^-^ B cell compartment, 437 upregulated and 190 downregulated DEGs (Fig. 3A, left) were identified. Similarly, in the CD19^+^CD138^+^ B cell subset, 427 DEGs were upregulated, and 274 DEGs were downregulated (Fig. 3A, right) (FDR *p*<0.05, >2-fold). Venn analysis revealed that both B cell subsets shared a core of DEGs (388 genes, 41.4%) when comparing cells recovered from pancreatic islets with those in the PLN, but also highlighted a substantial number of DEGs that distinguished these two B cell subsets (Fig. 3B, ESM Table 3). Volcano plots (Fig. 3C and D) for CD19^+^CD138^-^ and CD19^+^CD138^+^ respectively illustrate the genes that were up- (red dots) and down- (blue dots) regulated when islet B cell subsets were compared to PLN B cell subsets. Hierarchical clustering of the top 50 shared DEGs (Fig. 3E) in both B cell subsets (PLN vs pancreatic islets) demonstrated that the majority of genes were upregulated, including the key transcriptional regulator *Irf7* (Interferon Regulatory Factor 7) and *Tlr7*, the gene encoding Toll-like Receptor 7 (TLR7). Genes that were downregulated were associated with B cell activation, including *Cr2* (CD21), *Fcer2a* (CD23) and *Ciita* (Class II Major Histocompatibility Complex Transactivator). Finally, we measured the transcriptional distance between the B cell subsets in both tissues, revealing two discrete clusters, confirming the distinction between the populations found in PLN and pancreatic islets (Fig. 3F). Principal component analysis (PCA) highlighted the close relationships between the different CD19^+^ B cell subsets located within the respective tissues, particularly in the PLN. Taken together these data indicate that both B cell subsets are heavily influenced by their environment. They are transcriptionally similar when resident in the islets but are significantly different from their counterparts in the draining lymph node.

**Fig 3.**
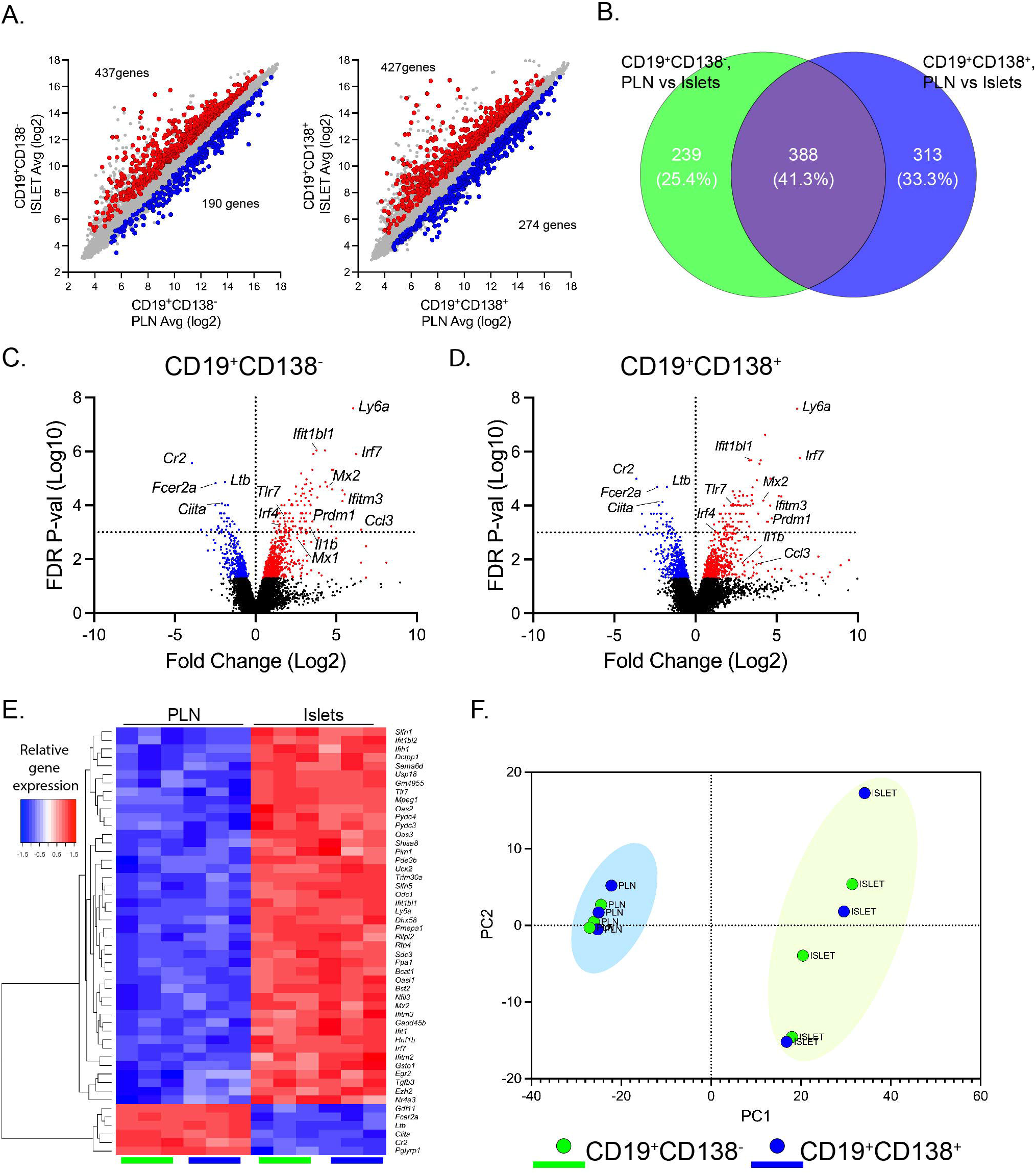
B cell populations are significantly modified by the inflamed pancreatic environment Pancreatic lymph nodes (PLN) and pancreatic islets from groups female NOD mice aged 16–20-week-old were taken (n=8) and CD19^+^CD138^-^ (green), CD19^+^CD138^+^ (blue) cells were FACS sorted for RNA isolation and gene array analysis. (A) Scatter plots to show differentially expressed upregulated (red) and downregulated (blue) genes between the PLN and the pancreatic islets from CD19^+^CD138^-^ and CD19^+^CD138^+^ B cell subsets. (B) Venn diagrams to show core shared genes and differences between both gene B cell gene sets upon location from PLN to pancreatic islets. (C, D) Volcano plots showing -Log10 FDR *p*-value (<0.01 indicated above dashed line) on the y-axis and Log2 fold change on the x-axis and annotated genes of interest. Differentially expressed genes between the PLN and pancreatic islets upregulated are shown in red and downregulated in blue (FDR<0.05). C: CD19^+^CD138^-^ D: CD19^+^CD138^+^. (E) Hierarchical clustering heatmap showing to top 50 most significant differentially expressed genes between the PLN and pancreatic islets for both B cell subsets (FDR<0.05, >2-fold). (F) Gene clustering of the differentially expressed genes (FDR<0.05, >2-fold) depicted by principal component analysis (PCA) plot.

### Islet-specific B cells are enriched in genes associated with antibody-secreting cells

Interestingly we noted that the genes *Prdm1* (Blimp1) and *Irf4*, both of which are key genes regulating plasma cell differentiation [32, 33], were upregulated in both islet CD19^+^ B cell subsets (in Figure 3). Therefore, we next compared the two gene sets from both B cell populations with previously published genes reportedly associated with the antibody-secreting cell (ASC) differentiation pathway, either as activated or repressed targets [22, 34] (Figure 4). Activated genes in the ASC list were compared with DEG genes in both gene sets, and Venn analysis revealed to our surprise, that CD19^+^CD138^-^ B cells, as well as CD19^+^CD138^+^, had a modest number of genes in common with genes activated in the ASC pathway (Fig. 4A). The majority of the shared DEG (between both B cell subsets and the ASC gene list) were activated (FDR *p*<0.05, >2-fold) and included the aforementioned *Prdm1* and *Irf4* (red arrows) (Fig. 4B). Additionally, *Ly6a* (Stem cell antigen-1 [Sca-1]) was highly upregulated in both gene sets (Fig. 3C, D) and encodes a surface antigen that has been used to improve plasma cell gating strategies in mice [35, 36]. Genes that were significantly altered in expression (FDR<0.01) and are included in an ASC gene signature [22] are listed for both CD19^+^CD138^-^ (green) and CD19^+^CD138^+^ (blue) B cells in Fig. 4. Expression values for *Prmd1, Irf4* and *Ly6a* genes in all samples are shown in ESM Fig. 3A and activation of the genes *Ly6a* and *Prdm1* in both B cell subsets in pancreatic islets, compared to the PLN, was confirmed by qPCR (ESM Fig. 3B). We next assessed the repressed genes identified in our B cell subsets which had also been previously identified as downregulated in ASC [22, 34]. Again, some genes were shared by the CD19^+^CD138^-^ and CD19^+^CD138^+^ populations, while others were exclusive to one or the other (FDR *p*<0.05, >2-fold), as demonstrated in a Venn diagram (Fig. 4C). Of the 92 shared genes, 23 were induced, including several IFN-induced genes (list of genes available in ESM table 4). However, key genes involved in antigen presentation, such as *Ciita* and *Cd40*, and B cell activation, e.g., *Fcer2a* (CD23), *Cr2* (CD21) and *Icosl* (red arrows), were repressed upon localisation of the cells to pancreatic islets (Fig. 4D).

**Fig 4.**
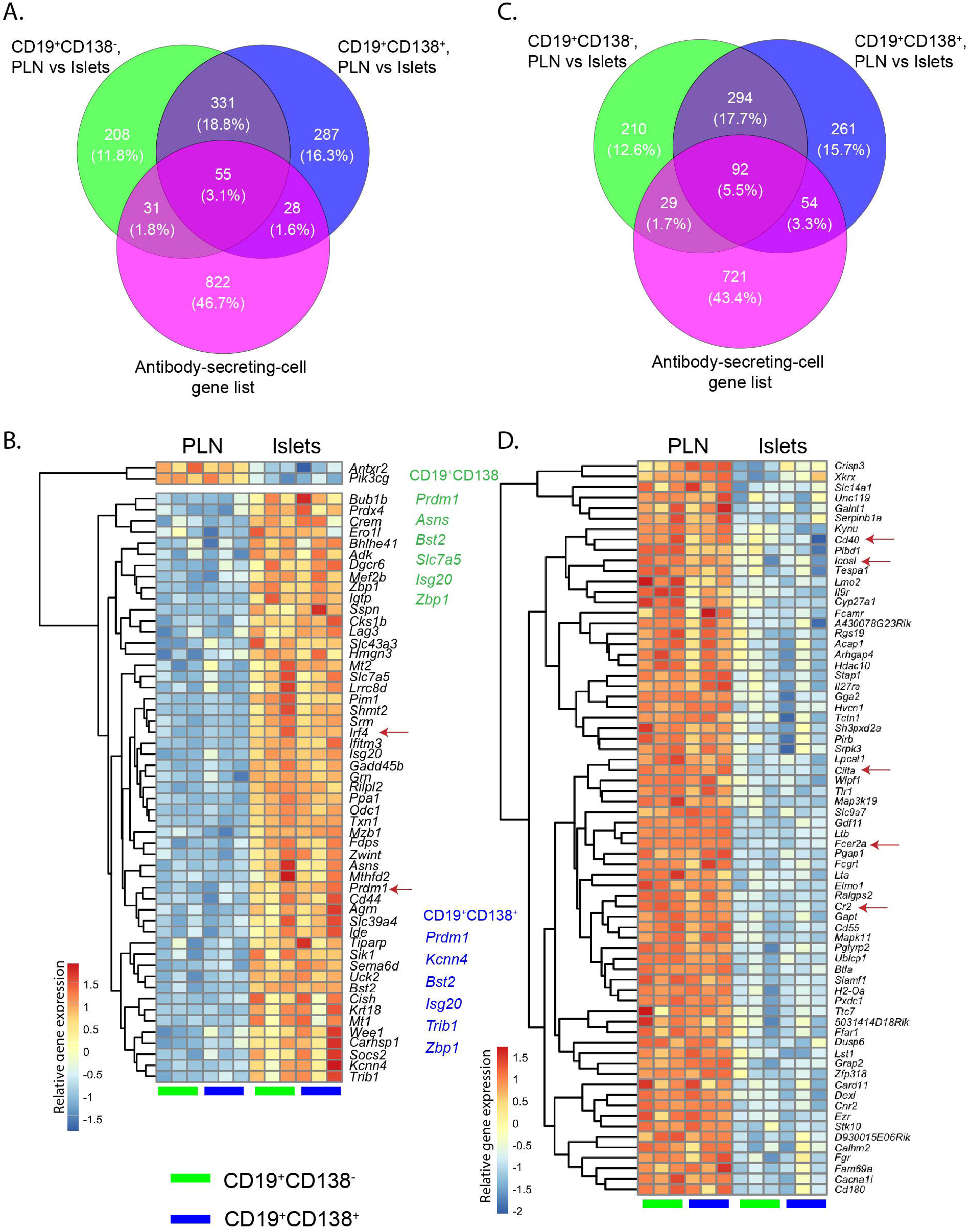
Islet-specific B cells are enriched in genes associated with antibody secreting cells Differentially expressed genes (FDR<0.05, >2-fold) from both B cell subsets from the PLN vs pancreatic islet comparison were assessed against published antibody secreting cell (ASC) genes, both overexpressed and repressed in the differentiation pathway. (A, C) Venn diagrams to show shared and non-shared genes significantly upregulated (A) and downregulated (C) between the PLN and the pancreatic islets in each cell subset and the ASC gene list. (B, D) Hierarchical clustering heatmap showing shared genes in pancreatic islets (compared to PLN) in both B cell subsets associated with ASC. B: Genes known to be overexpressed in ASC corresponding to B cell subsets. Genes listed are highly significant (FDR<0.01) and ASC signature genes in CD19^+^CD138^-^ (green) CD19^+^CD138^+^ (blue) B cell subsets. D: Genes known to be repressed in ASC corresponding to B cell subsets. Red arrows highlight key factors in B cell differentiation and function. Each column represents the relative gene expression from one experimental sample.

We next made efforts to understand if CD138^+^ B cells had more mature ASC-like changes upon translocation to the islets. Therefore, we interrogated the datasets to explore trends in the levels of expression of ASC-related genes, or exclusive genes in either B cell subset. The extent of the change for each B cell gene set that was upregulated and associated with ASC was plotted (ESM Fig. 4A). Key signature genes (such as *Prdm1* and *Irf4)* were induced to a similar extent (by comparison with their respective PLN subsets). A selection of genes unique to each cell subset was also identified. In the CD19^+^CD138^+^ B cell gene set, *Cd9* expression was increased by comparison with the CD19^+^CD138^-^ gene set (FDR <0.05 and fold change >2 cut off (ESM table 4)). Similarly, among the repressed genes, those such as *Bcl6* and *Il4ra, Ms4a1* (cell surface receptor CD20) were restricted to the CD19^+^CD138^+^ B cell subset. It was also noted that several of these genes were significantly changed in the CD19^+^CD138^-^ cells, but by <2-fold. ESM table 4 shows all differentially expressed ASC-related genes with an FDR <0.05, irrespective of the extent of the fold-change. Selectively expressed in CD19^+^CD138^-^ B cells were genes including *Alcam* (Leukocyte Cell Adhesion Molecule) and *Pycr1* (CD19^+^CD138^+^, FDR>0.1) (ESM Figure 4A). In contrast, CD19^+^CD138^+^ B cells selectively expressed *Tns3* and *Epcam* (CD19^+^CD138^-^, FDR>0.1).

To identify ASC genes that differed in their levels of expression, we selected genes with a >1.5-fold difference between the B cell gene sets (ESM Fig. 4B). Interestingly, using this criterion, the CD19^+^CD138^+^ B cell gene set had higher levels of expression of *Cd9* and *Cd44*, two genes which are associated with adhesion [37, 38]. *Mef2b*, a member of the *Bcl6* complex [39] was induced to a greater extent in the CD19^+^CD138^-^ subset. Other genes with the same pattern as *Mef2b* included *Pim1* and *Asns*, involved in cell cycle and survival [40, 41].

A substantial number of genes associated with the B cell activation pathway were repressed (ESM Table 4, FDR<0.05) and important genes such as *Cr2, Fcer2a* and *Ciita* were downregulated, in both cell populations, by comparison with those in the PLN (ESM Table 4). However, genes such as *Icosl, Il4ra, Bcl6, Ms4a1* and *Cd40* were repressed more substantially in the CD19^+^CD138^+^ B cell subset. Of note, *Pax5*, a gene directly repressed by *Prmd1*, was significantly downregulated by >-1.5-fold. To identify highly differential gene repression patterns, we again selected genes with >1.5-fold change difference (ESM Fig. 4C). *Il27ra*, a receptor for the cytokine IL-27 which supports germinal centre formation [42], was highly repressed in CD19^+^CD138^+^ compared to CD19^+^CD138^-^ cells. Again, a selection of genes was expressed uniquely in each B cell gene set (ESM Fig.4C). In CD19^+^CD138^-^ B cells, very few uniquely altered genes were identified, but these included *Birc3*, a regulator of the non-canonical NF-κB complex [43]. By contrast, in CD19^+^CD138^+^ cells, several genes were selectively repressed, including, for example, *Map4k2* (encoding MAP kinase 2; also known as germinal centre kinase). These data suggest that CD138^+^ B cells have a more substantial number of genes repressed upon islet translocation, compared to CD138^-^ B cells, but they do not show expression of a single group of genes characteristic of a more mature ASC cell. Overall, these data suggest that both B cell subsets found in pancreatic islets arrive in an inflamed environment, which favours the induction of key antibody-secreting cell-associated genes.

### Identifying gene expression differences in CD19^+^CD138^+^ B cells in pancreatic islets

A further goal of our gene profiling experiments was to identify any additional transcriptional changes in the CD19^+^CD138^+^ B cell subset (enriched in autoreactive B cells), therefore, we extended our investigation using gene ontology (GO) analysis. First, the functional annotation tool DAVID [44] was employed (ESM Figure 5), and it was found that GO term ‘cell cycle’ (GO0007049) was significantly enriched in the CD19^+^CD138^+^ B cell gene set (FDR *p*<0.002, enrichment score 2.49) but not in CD19^+^CD138^-^ B cells (FDR *p*<0.08, enrichment score 2.02) (data not shown).

**Fig 5.**
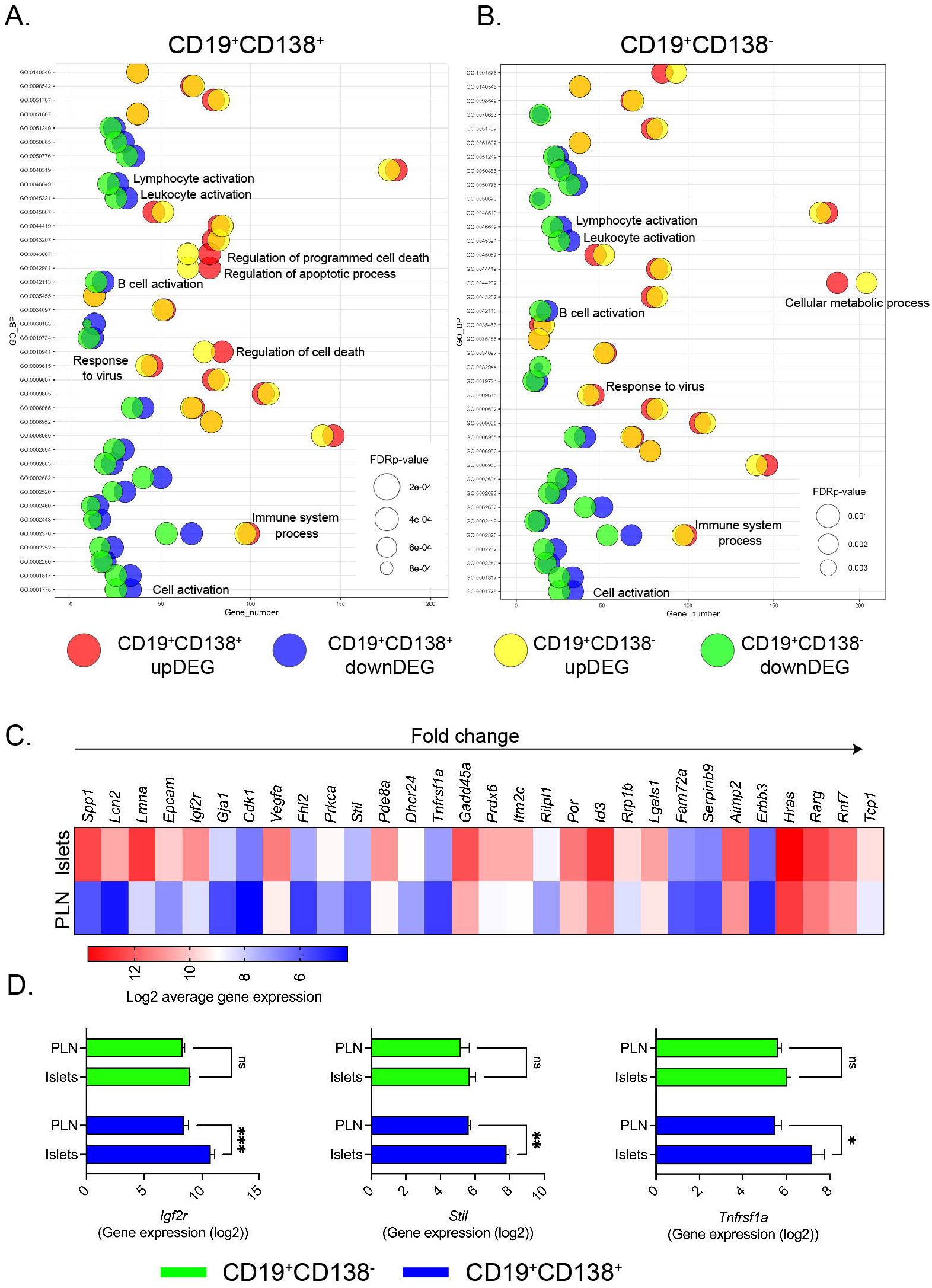
CD19^+^CD138^+^ B cells in the pancreas overexpress growth factor and late cell cycle genes Genes either up-or-downregulated in each gene set (FDR *p*<0.05, >2-fold change) were used to test for over-represented biological processes using PANTHER Classification System. (A, B) Bubble plots demonstrating top 20 overrepresented gene ontology (GO) terms in each B cell subset. GO terms are plotted against related gene number identified in either B cell subset. FDR *p*-value is depicted by size of the bubble. (A) GO terms found enriched in CD19^+^CD138^+^ B cell gene set (B) GO terms found enriched in CD19^+^CD138^-^ B cell gene set. CD19^+^CD138^+^ upregulated DEG (CD19^+^CD138^+^ upDEG; red), CD19^+^CD138^-^ upregulated DEG (CD19^+^CD138 upDEG; yellow), CD19^+^CD138^+^ downregulated DEG (CD19^+^CD138^-^ downDEG; blue), CD19^+^CD138^-^ downregulated DEG (CD19^+^CD138^+^ downDEG; green). DEG: differentially expressed genes. (C) Heatmap showing average expression of upregulated DEG (FDR *p*<0.05, >2-fold change) in CD19^+^CD138^+^ B cell gene set (PLN and pancreatic islets) identified in enriched GO terms related to ‘Cell death’ and the ‘Apoptotic process’. (D) Average (mean ± SEM) gene expression of *Igf2r, Stil, Tnfrsf1a* in B cell subsets for both PLN and pancreatic islets. ***<0.001, **<0.01, *<0.05, two-way ANOVA with a Bonferroni’s multiple comparison test.

We next used the PANTHER Classification System and performed a more “in-depth” gene ontology analysis on the DEGs in both CD19^+^CD138^+^ and CD19^+^CD138^-^ B cells (Figure 5). Genes either up-or-downregulated in each set (FDR *p*<0.05, >2-fold change) were used to test for over-represented biological processes. The bubble plot in Fig. 5A shows the most significant biological processes from DEGs either upregulated (top 20 GO terms) (red dots) or downregulated (top 20 GO terms) (blue dots) in CD19^+^CD138^+^ B cells. Gene number is plotted against GO terms, while FDR *p*-value is indicated by size of the bubble. GO terms in the double-positive B cells were compared with gene numbers and corrected *p*-value in the up (yellow) and down-regulated (green) DEGs in CD19^+^CD138^-^ B cells. Fig. 5B shows a bubble plot of the top GO terms for the CD19^+^CD138^-^ B cells, with gene number and corrected *p*-value compared with the double-positive B cells. A full description of GO terms is provided in ESM Figure 6.

**Fig 6.**
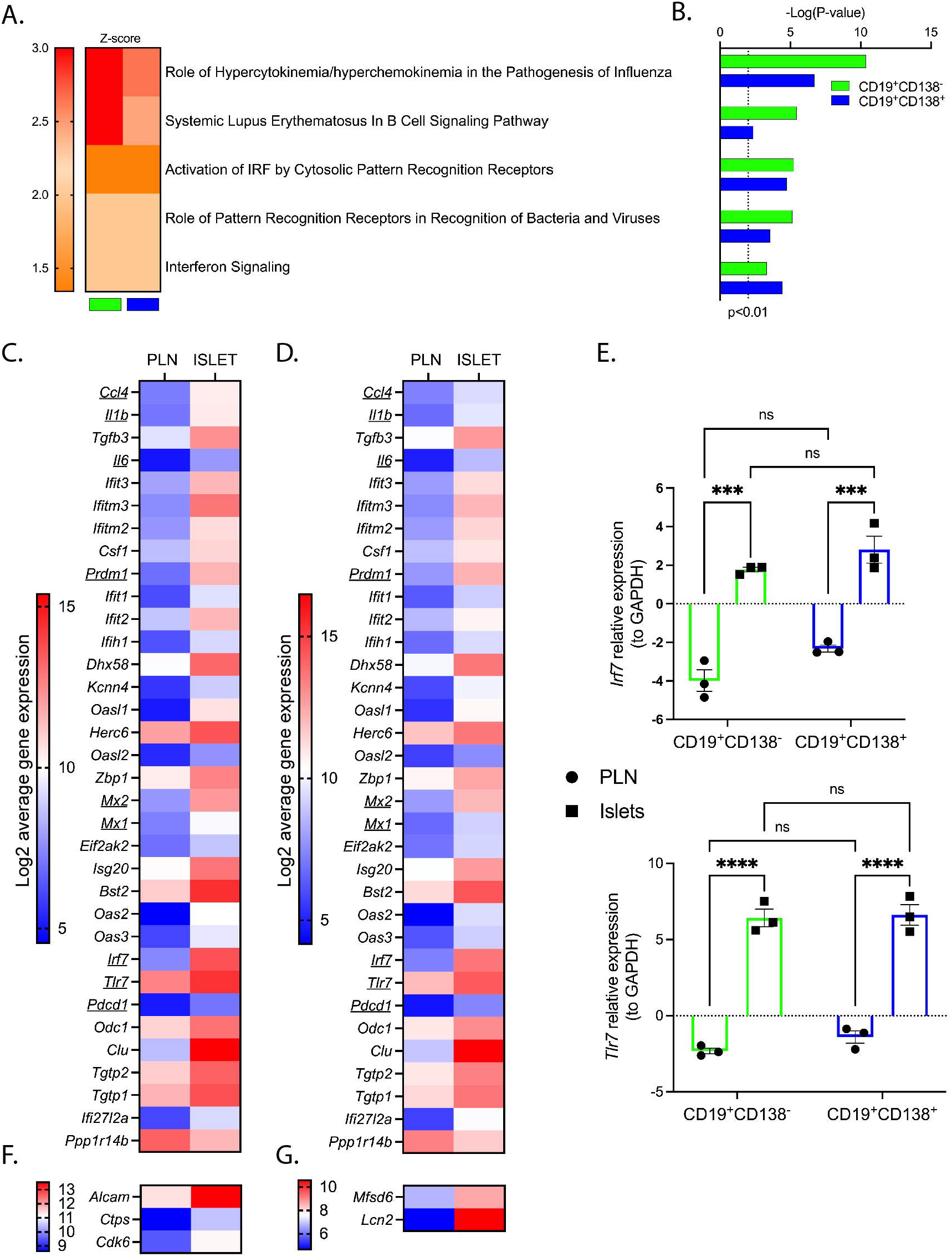
CD19^+^ B cell subsets acquire an innate immune signature upon location to pancreatic islets. Enriched gene canonical pathways and networks in B cell subsets (CD19^+^CD138^-^; green, CD19^+^CD138^+^; blue) in pancreatic islets using Ingenuity Pathway Analysis software (FDR <0.05, 1.5-fold-change). (A, B) Canonical pathways in B cell subsets displaying significant activation (orange-red) denoted by A: z-score B: corresponding corrected *p*-value, dotted line represents *p*<0.01. (C, D) Expression profile of innate or interferon related signature genes shared between subsets. Underlining highlights genes of interest. C: CD19^+^CD138^-^, D: CD19^+^CD138^+^ (E) qPCR of *Irf7* and *Tlr7* genes in B cell subsets located in the PLN and pancreatic islets. Expression was measured to housekeeping gene GAPDH. ***<0.001, ****<0.0001, two-way ANOVA with a Bonferroni’s multiple comparison test. (F, G) Expression profile of innate or interferon related signature genes not shared between subsets. F: CD19^+^CD138^-^G: CD19^+^CD138^+^.

Many of the top GO terms overlapped (both in gene number and corrected *p*-value), including ‘immune system process’ (GO0002376) and ‘response to virus’ (GO0009615), although some important differences were observed. The CD19^+^CD138^-^ B cell subset had a higher enrichment of upregulated genes associated with metabolic processes (GO0044237). While in the CD19^+^CD138^+^ B cells, genes relating to B cell activation were downregulated while there were increases in upregulated genes involved with ‘programmed cell death’ (GO0043067) or the ‘apoptotic process’ (GO0042981). We, therefore, studied the genes associated with cell death processes and selected 30 genes that were significantly expressed in CD19^+^CD138^+^ (compared to CD19^+^CD138^-^, FDR>0.05) (Fig. 5C). The top 5 genes identified were *Spp1* (Osteopontin), *Lcn2* (Lipocalin 2), *Lmna* (Laminin A), *Epcam* (EpCAM, adhesion molecule) and *Igfr2* (receptor for insulin growth-like factor 2). Other genes also featured, such as *Cdk1* (Cyclin dependent kinase 1) and *Gadd45a*, both cell cycle genes.

Genes such as *Spp1* and *Lmna* yielded a corrected *p*-value <0.08 in the CD19^+^CD138^-^ gene set (data not shown). Therefore, we eliminated genes from the list in Fig. 5C that were expressed in the single positive B cell gene list and had a p<0.05 (not corrected). This left 8 genes (*Igfr2, Stil, Tnfrsf1a, Fam72a, Serpinb9, Erbb3, Rnf7* and *Tcp1*) identified as being expressed exclusively in CD19^+^CD138^+^ B cells. The values for the top 3 DEG genes having significant differences between CD19^+^CD138^+^ B cells in the PLN vs islets are presented in Fig 5D. *Igf2r* encodes the receptor that binds the growth factor IGF2, which has a role in cell proliferation [45] and is also overexpressed in mature bone-marrow plasma cells [46]. Expression of *Igfr2* could play an anti-apoptotic role in CD19^+^CD138^+^ B cells as knockdown of this receptor in cervical cancer cells induces apoptosis [47]. Similarly, *Stil* (STIL) has an important role in the cell cycle, specifically centriole assembly [48]. Interestingly overexpression of *Tnfrsf1a* (TNF Receptor 1[TNFR1]), a growth factor for B cells [49] and a cytokine implicated in the pathogenesis of type 1 diabetes, could indicate a heightened response to any locally available TNFα in pancreatic islets. Altogether, these data suggest that CD19^+^CD138^+^ B cells have a subtle but clear difference in their gene profile when compared to CD19^+^CD138^-^ B cells, upon trafficking to the pancreas. Upon arrival in the islet vicinity, double-positive B cells appear to overexpress genes associated with adhesion, growth factors and later stages of the cell cycle.

### CD19^+^ B cell subsets acquire an innate immune signature upon translocation to pancreatic islets

We next explored enriched gene regulatory networks in the islet-specific B cells using our dataset and IPA (Ingenuity Pathway Analysis) (Figure 6). This revealed the top 5 canonical pathways that displayed significant increases in activation (corrected *p*<0.01). The activation status of each canonical pathway, for each CD19^+^ B cell subset, is denoted by the z-score (orange = activated) in the heatmap shown in Figure 6A, with corresponding *p* values for each pathway in Fig. 6B. Top pathways activated in both gene sets were ‘Role of Hypercytokinemia/hyperchemokinemia in the Pathogenesis of Influenza’ and ‘Systemic Lupus Erythematosus In B Cell Signalling Pathway’. Other activated pathways included activation of PRR (Pattern Recognition Receptors), indicating that B cells were enriched for expression of interferon-induced genes. We used the activated IPA pathways and DAVID GO analysis (ESM Fig. 5) to identify the key signature genes upregulated by CD19^+^CD138^-^ (Fig. 6C) and CD19^+^CD138^+^ (Fig. 6D) B cells after translocation to the pancreatic during established insulitis. The majority of genes were significantly upregulated in both B cell subsets, including cytokines *Il6, Il1b* and *Ccl4* (MIP-1β), transcription factors *Irf7* and *Prdm1* and key anti-viral proteins *Mx1* and *Mx2*. Other overexpressed genes included *Tlr7* and *Pdcd1* (Programmed Cell Death 1[PD-1]), and we were able to confirm the upregulation of *Tlr7* and *Irf7* in both islet-localised B cell subsets by qPCR (Fig. 6E). Three DEG were upregulated in the CD19^+^CD138^-^ gene set but not in that from CD19^+^CD138^+^ cells (FDR>0.1), including *Alcam* (also identified above) and the early cell cycle gene *Cdk6* (Fig. 6F). Specific to the CD19^+^CD138^+^ B cells was upregulation of viral response genes *Lcn2* and *Mfsd6*, the latter involved in MHC I antigen presentation (Fig. 6G).

### TLR7 protein expression is evident in islet-specific B cells

Finally, we sought to confirm the expression of the TLR7 protein in the pancreas of NOD mice that had developed diabetes (Figure 7). TLR7 expression was visible in insulin-expressing beta cells in the few remaining insulin-containing islets (Fig. 7A) and many B cells with florid CD20 expression (Fig. 7B). Double positive (CD20^+^TLR7^+^) B cells were also observed in or around insulin-deficient islets (Fig. 7C). Furthermore, in the remaining islet structures and immune cell clusters, we observed both TLR7^+^ CD20^+^ cells and TLR7^+^ CD20^-^ cells in close proximity (Fig. 7D). We also examined TLR7 expression on islet B cells using flow cytometry (Fig. 7E-G), enabling demarcation of our CD138 B cell subsets in both the PLN and pancreatic islets (Fig. 7E). TLR7 expression was significantly increased on both islet CD19^+^ B cell subsets compared to their PLN counterparts, corroborating our gene expression observations (Fig. 7F, G). To note CD19^-^CD138^+^ cells (grey shaded gate) showed little TLR7 expression (data not shown). However, we did observe a higher level of expression of TLR7 on CD19^+^CD138^+^ B cells compared to the CD19^+^CD138^-^ subset in pancreatic islets, and this was not observed in the PLN (Fig. 7G).

**Fig 7.**
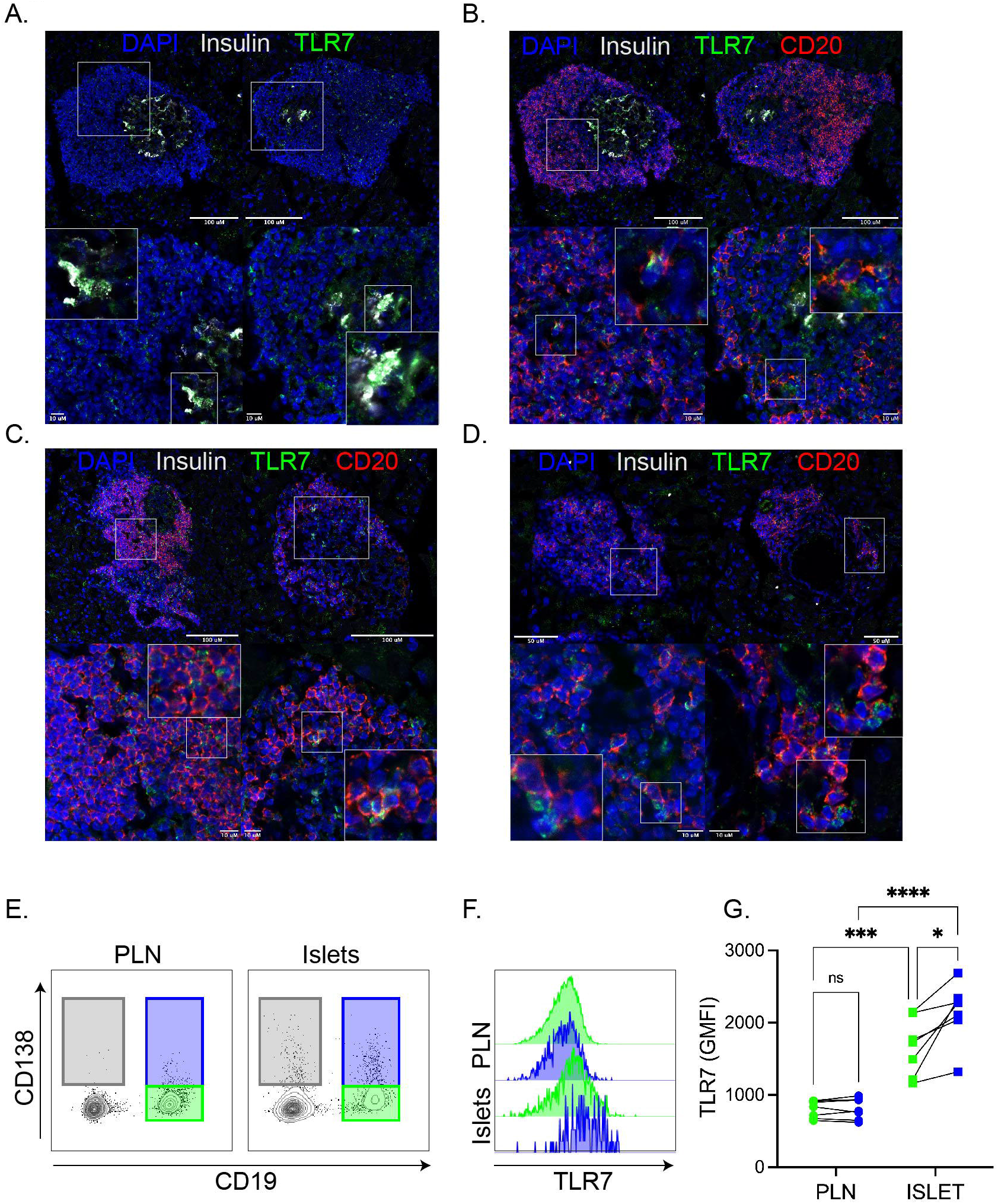
Expression of TLR7 protein in pancreatic islets in NOD mice. (A-D) Representative confocal microscopy images of pancreatic sections from diabetic NOD female mice (blood glucose >13.9mmol/L) stained with insulin (grey), CD20 (red), TLR7 (green) and nuclear DAPI counterstain showing in remaining insulin containing islets (A, B), insulin-expressing beta cells positive for TLR7 (A) CD20^+^TLR7^+^ B cells in remaining insulin containing islets (B). CD20^+^TLR7^+^ B cells in insulin deficient islets (C). CD20^+^TLR7^+^ B cells in close contact with single positive TLR7^+^ cells (D). Images represent pancreatic islets or immune cell clusters in 3 individual mice. (E-G) Pancreatic lymph nodes and pancreatic islets from female NOD mice aged 16-20 weeks old were taken before flow cytometric analysis on (E) CD19^+^CD138^-^ (green gate), CD19^+^CD138^+^ (blue gate) B cells and CD19^-^CD138^+^ cells (grey gate). (F) Histogram showing TLR7 expression on CD19^+^ B cell subsets (G) Summary graph of TLR7 (geometric mean fluorescent intensity, GMFI) expression. Each dot represents one individual mouse over two independent experiments. ***<0.001, ****<0.0001, two-way ANOVA with a Bonferroni’s multiple comparison test.

## Discussion

We show that, in NOD mice with established insulitis, the gene expression profiles of B cells are influenced dramatically by the pancreatic environment, with inflammation favouring the upregulation of IFN-associated genes and the induction of genes promoting plasma cell differentiation. Gene expression analysis revealed a clear distinction between the populations of B cells found in the pancreatic lymph nodes vs those infiltrating the islets during disease development. Activated pathways were associated with innate immune signalling and included the upregulation of genes such as *Irf7* and the toll-like receptor *Tlr7* alongside multiple pro-inflammatory cytokines such as those encoded by *Il6* (IL-6), *IL1b* (IL-1β) and *Ccl4* (MIP-1β). Interestingly, repressed genes were associated with antigen presentation and B cell activation, including *Ciita* and *Cd40* suggesting that islet-specific B cells may primarily facilitate local beta cell damage by producing pro-inflammatory cytokines.

Our approach also highlighted a population of CD138^int^ cells enriched (compared to secondary lymphoid organs) in pancreatic islets of NOD mice, that we have previously described as a heterogeneous population of plasmablast or plasma-like cells with downregulated CD19 and IgD but encompassing a population of insulin-specific B cells [11]. These CD138^+^ cells remain in the minority and are not selectively expanded in the pancreas during diabetes development [12] (data not shown). Gene expression analysis revealed a large number of genes that are differentially expressed compared to those in CD19^+^ B cells, including key innate lymphocyte genes and plasma cell-related genes. Recent work shows that CD138 is a marker for NKT17 cells [23], suggesting that in pancreatic islets, such cells might be present during diabetes development. In support of this, the *Il17re* gene was activated to a greater extent in the CD138^+^CD19^-^ population, by comparison with CD19^+^ B cells. We also discovered that the population of CD138^+^ cells is enriched in differentiated plasma cells since *Jchain* was highly upregulated, and a small number of IgA^+^ plasma cells were detected. Notably, evidence in other autoimmune diseases such as multiple sclerosis (MS), implies that IgA^+^ B cells are enriched in inflammatory lesions in the central nervous system (CNS) in both humans [50] and mice [51], with a role in attenuating disease via IL-10 production. The roles, significance and origin of IgA^+^ plasma cells in the pancreas are currently unknown and are worthy of further investigation.

Analysis of the gene expression profile of CD19^+^ B cells in the pancreas, as compared to that of equivalent cells in the lymph node, revealed an induction of both IFN and plasma cell-related genes. It is well documented that Blimp-1 and IRF4 are key transcription factors regulating plasma cell differentiation [30, 52] and that the expression of Blimp-1 leads to repression of the B cell commitment gene *Pax5*, among others [34]. Directly targeted by Blimp-1 expression is the MHC II regulating gene *Ciita* [32] which we show is highly repressed in B cells that have migrated to pancreatic islets. Repression of Pax5 is required for the initiation of ASC development [53] and is repressed in B cells present in pancreas (FDR<0.05, FC>1.5), however no difference was seen in the expression of the transcription factor, XBP-1 (X-box binding protein-1). XBP-1 is a target of Pax5 and acts downstream of Blimp-1 to regulate the UPR (unfolded protein response) [31, 54], a process essential for the secretion of immunoglobulins. These results imply that islet-specific CD19^+^ B cells upregulate genes associated with antibody secretion but are not actively engaged in secreting immunoglobulins. Based only on gene expression profiles, it is difficult to discern whether pancreatic CD19^+^ B cells are at the stage of pre-plasmablasts or whether they may still be precursors, as many genes in this pathway are transitional or are expressed continuously [28]. Functional assays and assessment of protein expression using CD19^+^ B cells from both the lymph node and pancreas may provide more definitive evidence.

Plasma cell-associated genes such as Blimp-1 and IRF4 can be activated by factors other than BCR signalling, such as IL-21 and IL-6 [55] or IFNα and IL-6 [56], cytokines variously implicated in the pathogenesis of type 1 diabetes. IL-21 can directly induce *Prdm1* gene expression, and this requires both STAT3 and IRF4 expression [57], which were each induced in islet-localised B cells in this study. IFNα and IL-6, produced by plasmacytoid DCs (pDCs), can cause CD40-activated B cells to differentiate into ASCs [56]. Crosstalk between IFNα-producing pDCs and B-1a cells in the pancreatic islets during the early stages of autoimmune diabetes is crucial for initiation of disease [7], and our data demonstrate that B cells are influenced by, and acquire, this innate signature during the establishment of insulitis. It is well documented that type 1 interferon is a major player in the pathogenesis of type 1 diabetes, since IFNα is expressed by the beta cells of patients with type 1 diabetes [58], and IFN-associated genes are overexpressed in islets of newly diagnosed individuals with type 1 diabetes [59]. Type 1 IFNs enhance the expression of, and response to, TLR7 [60, 61], a gene product highly expressed in the B cells found in pancreatic islets of NOD mice. Furthermore, in combination with IFNα, TLR7 activation augments IL-6 production and isotype switching in B cells [60]. In NOD mice, TLR7 deficiency delays and reduces the development of autoimmune diabetes and alters the functional responses of B cells [62]. Therefore, it is likely that the heightened expression of IFNα enhances TLR7 expression in B cells situated in the pancreas, and that this, in turn, synergistically amplifies IL-6 and pro-inflammatory cytokine production. The status of TLR7 in those CD20^+^ B cells found in association with the pancreatic islets of human subjects with type 1 diabetes is currently unknown, but given that an elevated proportion of infiltrating B cells correlates with earlier diagnosis and more rapidly progressive disease [6], targeting of TLR7 could be of interest therapeutically.

An important finding from our work is the heightened expression of the TLR7 protein in CD19^+^CD138^+^ islet B cells, which encompass a substantial proportion of the B cell population in the pancreas during the development and onset of diabetes in NOD mice. Previously, the CD138^int^ B cell population has been described as a pre-plasma cell (pre-PC) subset that are localised mainly in the follicular region of the spleen [63, 64] and which do not require Blimp-1 (*Prdm1*) expression for differentiation [30]. Importantly, a substantial number of CD138^int^ B cells are autoreactive [63], and we have observed that the population of insulin-specific B cells in NOD mice is enriched within the CD138^int^ subset [11]. Understanding the role of both CD138^-/+^ B cells and their relationship in the pancreatic tissue may have important consequences for B cell-targeted immunotherapy.

Our gene array data revealed little difference in the CD19^+^ B cell subsets when localised within the tissue (either lymph node or pancreas), as these subsets displayed a similar gene expression profile. However, distinct signatures were found when comparing the equivalent B cell subsets in the pancreatic lymph node and pancreas, notably in the expression of selected genes such as *Igf2r*. In mice lacking the growth factor *Igf2*, DCs (dendritic cells) have a reduced propensity to activate T cells [65], and overexpression of IGF2 in beta cells enhances their susceptibility to damage and stress [66]. Interestingly, antigen-specific B cells express high levels of IGF2R in mice treated with a regimen of OVA (Ovalbumin), and after ligation by IGF2, enhanced proliferation and IL-10 production was observed [67]. Further studies are required to fully understand if CD138^+^ B cells fulfil a specific functional role or if they are more responsive to TLR7 ligands. However, it is worth considering that the two subsets of CD19^+^ B cells are at different stages of differentiation or activation when harvested from the tissue, particularly as CD138 (syndecan-1) can be shed from the cell surface after ligand engagement [68].

Further studies using unbiased single-cell RNA sequencing have recently been performed in the pancreas of NOD mice during diabetes development [69], and application of an approach such as this may shed further light on the heterogeneity among these specific B cell subsets.

A further limitation of the present study is that we do not address B cell-specificity, which, if understood more fully, may help to highlight key differences in functionality of the CD138^+^ B cell subsets. Despite this, our observations provide new and important findings indicating how B cells may contribute to local beta cell damage and perpetuate tissue inflammation. Taken together, our study provides novel insights into potential therapeutic avenues which may be effective in individuals with type 1 diabetes.

## Supporting information

ESM Figures

## Data availability

The datasets generated and/or analysed during the current study are available from the corresponding author on reasonable request.

## Funding

This work was funded by Diabetes UK (19/0006032). JB is supported by an Independent Fellowship funded by Research England’s Expanding Excellence in England (E3) fund via EXCEED.

## Duality of interest

The authors declare that there is no duality of interest associated with this manuscript.

## Contribution statement

JB designed the experiments, acquired and analysed the data and wrote the manuscript. JD and JH contributed to experimental procedures. JP, PL, SR, NGM and FSW contributed to interpreting the results and revised the manuscript. All authors have reviewed and approved the manuscript. JB conceived the project and is the guarantor of this work.

## Acknowledgements

We thank Central Biotechnology Services at Cardiff University and the Histology Services Unit at the University of Bristol.

## Figure legends

**ESM Fig 1** Comparison of CD19^+^CD138^-^ and CD19^+^CD138^+^ B cells. (A) B cells from NOD splenocytes were stained for B cell markers including CD138 to show CD138^+^ B cells are enriched in the mature B cell population (B) Gating strategy for FACS sorting B cells in the PLN (top) and pancreatic islets (bottom) (C) Violin plots for the relative gene expression acquired from the ImmGen database for B cell populations. Genes are related to expression differences between CD19^+^CD138^-^ and CD19^+^CD138^+^ B cells in the PLN (left) and the pancreatic islets (right) shown in Figure 2A.

**ESM Fig 2** The CD19^-^CD138^+^ subset contain innate lymphocyte population. Gene expression related to Figure 2. (A, B) Hierarchical clustering heatmaps to show the most significant (FDR *p*<0.05, >2-fold) change genes differentially expressed when comparing CD19^+^CD138^-^ B cells to CD19^-^CD138^+^ subsets in both (A) PLN and (B) pancreatic islets. (C) *Jchain* gene expression from ImmGen database demonstrating exclusive expression in plasma cells (B. PC. Sp).

**ESM Fig 3** Pancreatic islet specific B cells upregulate antibody secreting genes. Related to Figure 4. (A)Average (mean ± SEM) gene expression of *Prmd1, Irf4* and *Ly6a* in CD19^+^CD138^-^ (green bars) and CD19^+^CD138^+^ (blue bars) B cell subsets for both PLN and pancreatic islets (B) qPCR of *Ly6a* and *Prdm1* genes in B cell subsets located in the PLN and pancreatic islets. Expression was measured to housekeeping gene GAPDH. *<0.05, ****<0.0001, ns: not significant, two-way ANOVA with a Bonferroni’s multiple comparison test.

**ESM Fig 4** Islet-specific B cells are enriched in genes associated with antibody secreting cells. Bar graphs showing Log2 fold change (left y-axis) and FDR *p*-value (right y-axis) for CD19^+^CD138^-^ (green bars) and CD19^+^CD138^+^ (blue bars) B cell gene sets (PLN vs pancreatic islets). Dotted line indicates FDR *p*-value 0.05. Genes not expressed in B cell subsets (single bars) had an FDR *p*-value>0.1. (A) Activated antibody secreting cell signature genes (B, C) Genes with a dissimilar fold change (>1.5-fold change difference) between CD19^+^CD138^-^ and CD19^+^CD138^+^ that were activated (B) or repressed (C).

**ESM Fig 5** Gene ontology using DAVID for both B cell subsets. Differentially expressed genes upregulated (FDR *p*<0.05, >2-fold change) were uploaded to the functional annotation tool DAVID. Bar graphs show corrected *p*-value for enriched gene ontology terms clustered by DAVID in CD19^+^CD138^-^ cells and CD19^+^CD138^+^ B cells.

**ESM Fig 6** List of Gene Ontology (GO) terms. Related to Figure 5.

**ESM Table 1** CD19^+^CD138^-^ B cells vs CD19^-^CD138^+^ in the PLN (Figure 2, ESM Figure 2)

**ESM Table 2** CD19^+^CD138^-^ B cells vs CD19^-^CD138^+^ in the pancreatic islets (Figure 2, ESM Figure 2).

**ESM Table 3** Shared and distinct genes in CD19^+^CD138^-^ (PLN vs pancreatic islets) and CD19^+^CD138^+^ (PLN vs pancreatic islets) (Figure 3).

**ESM Table 4** Antibody secreting cell genes expressed in B cell gene sets FDR<0.05 (Figure 4, ESM Figure 4).

## Notes

### Competing Interest Statement

The authors have declared no competing interest.

