## Supplementary material for "Gene expression profiling reveals B cells are highly educated by the pancreatic environment during autoimmune diabetes": ESM Figures

ESM Figure 1.

A.

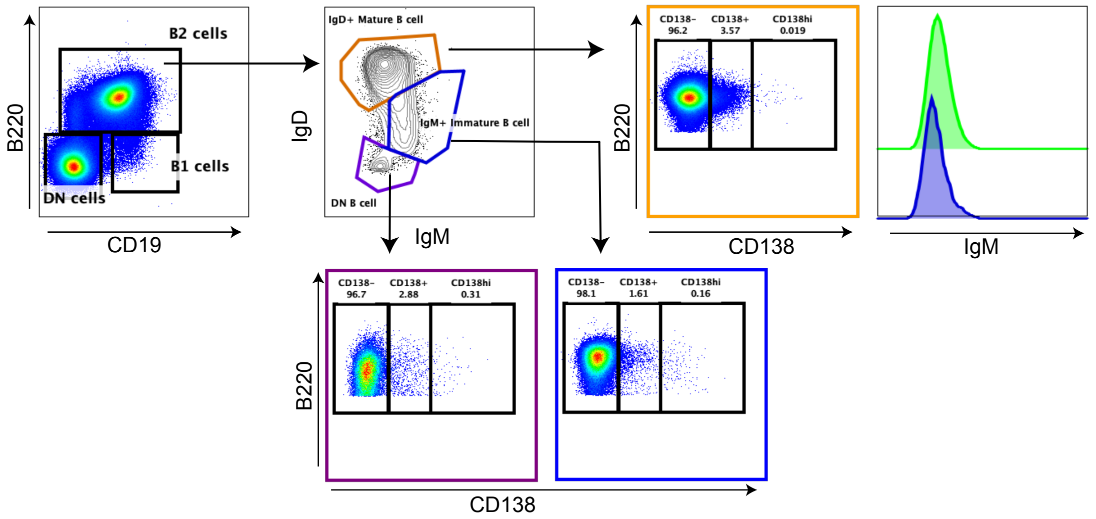


B.

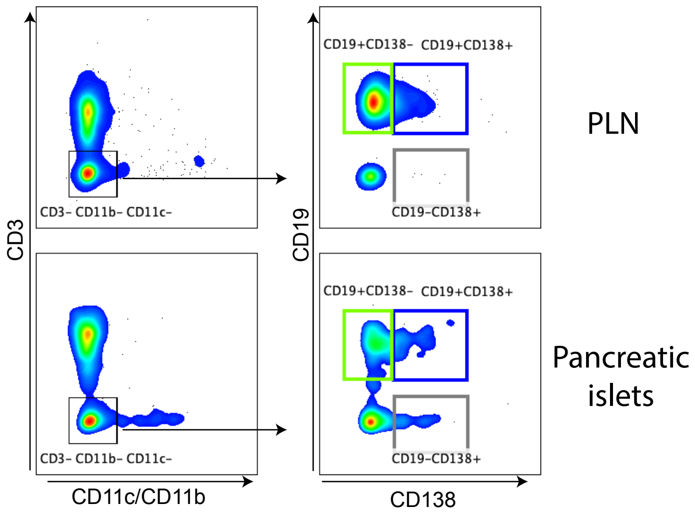


C.

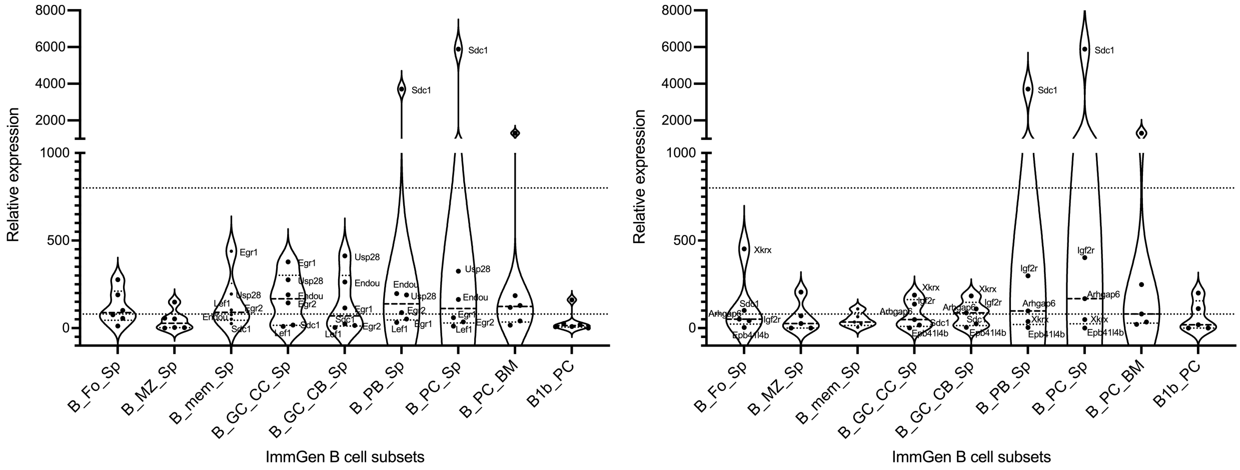

ESM Figure 2.

A.

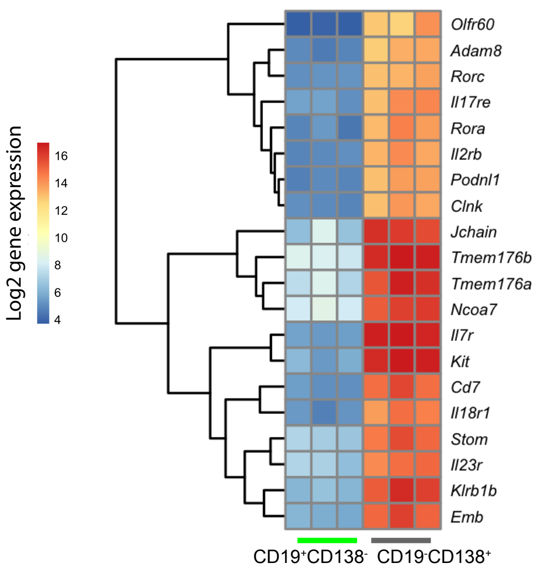


B.

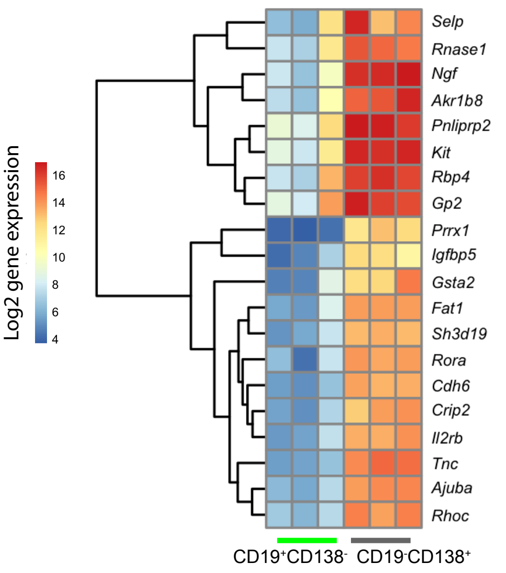


C.

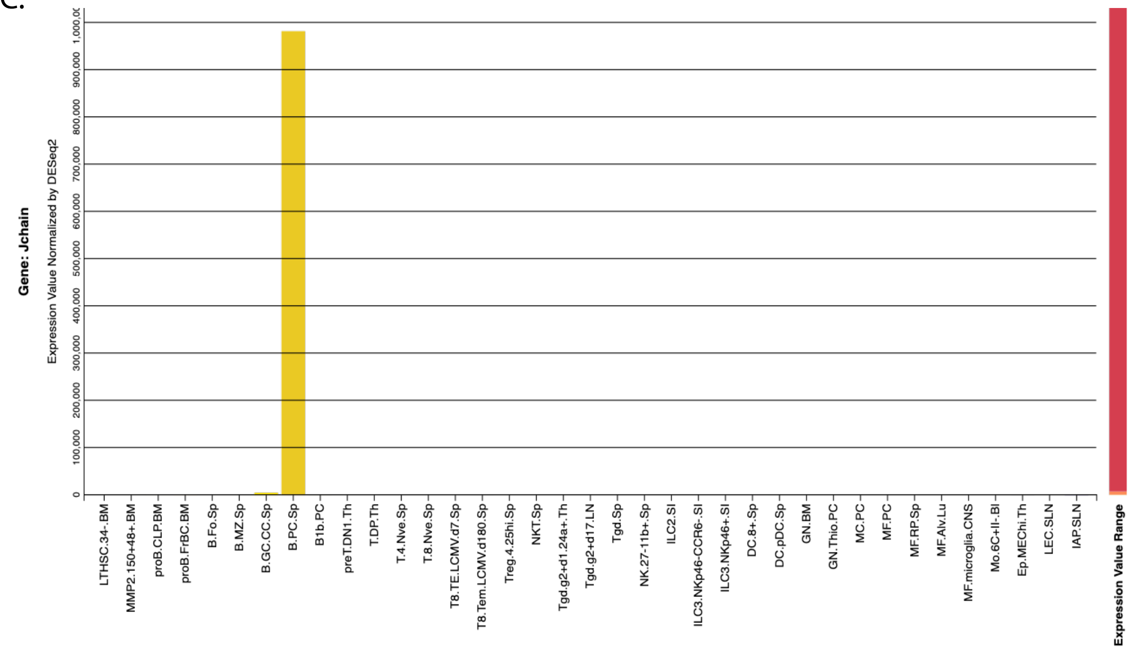

### ESM Figure 3.

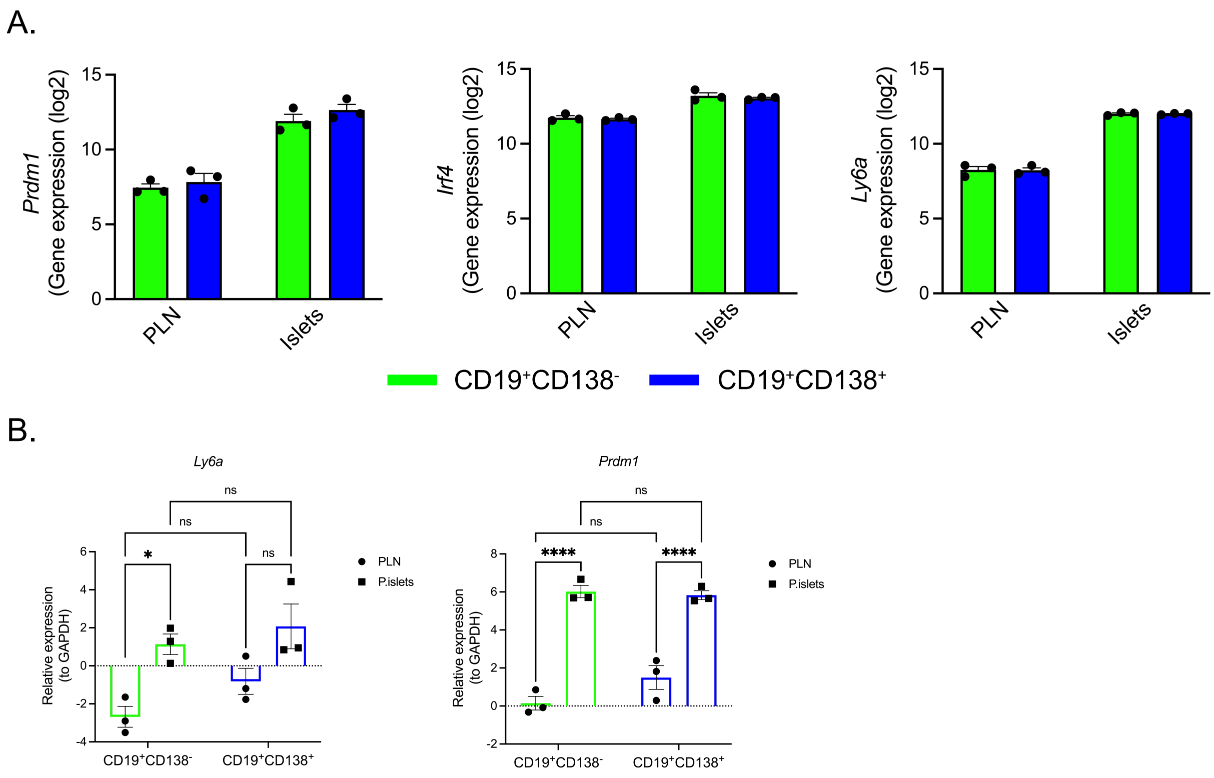

A.
Prdm1 (Gene expression (log2))
Irf4 (Gene expression (log2))
Ly6a (Gene expression (log2))
PLN
Islets
CD19+CD138-
CD19+CD138+
B.
Ly6a
Prdm1
Relative expression (to GAPDH)
ns
*
****
PLN
P.islets
CD19+CD138-
CD19+CD138+

ESM Figure 4.

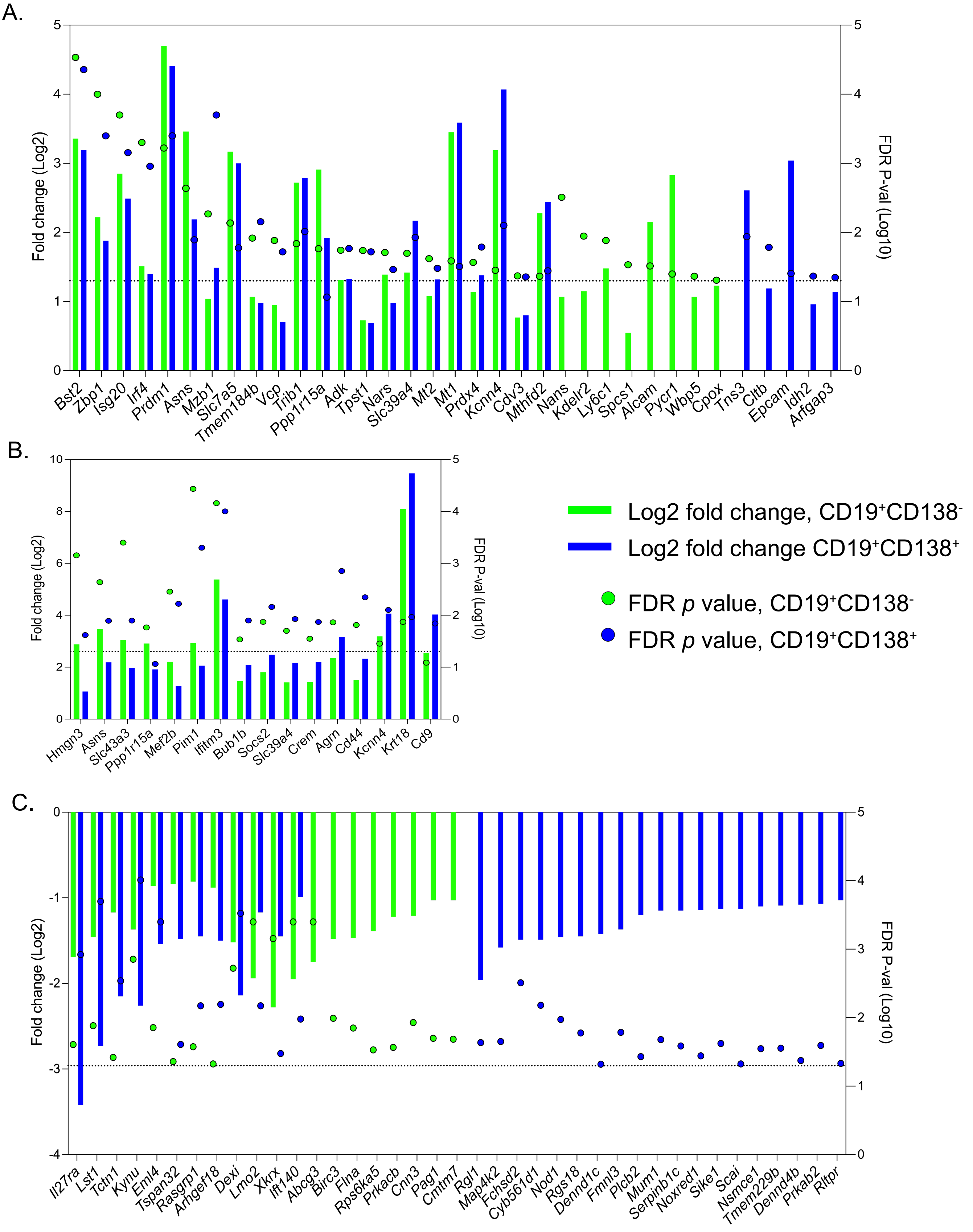
A.
Fold change (Log2)
FDR P-val (Log10)
Bst2
Zbp1
Isg20
Irf4
Prdm1
Asns
Mzb1
Slc7a5
Tmem184b
Vcp
Trib1
Ppp1r15a
Adk
Tpst1
Nars
Slc39a4
Mt2
Mt1
Prdx4
Kcnn4
Cdv3
Mthfd2
Nans
Kdelr2
Ly6c1
Spcs1
Alcam
Pycr1
Wbp5
Cpox
Tns3
Cltb
Epcam
Idh2
Arfgap3
B.
Fold change (Log2)
FDR P-val (Log10)
Hmgn3
Asns
Slc43a3
Ppp1r15a
Mef2b
Pim1
Ifitm3
Bub1b
Socs2
Slc39a4
Crem
Agrn
Cd44
Kcnn4
Krt18
Cd9
Log2 fold change, CD19+CD138-
Log2 fold change CD19+CD138+
FDR p value, CD19+CD138-
FDR p value, CD19+CD138+
C.
Fold change (Log2)
FDR P-val (Log10)
Il27ra
Lst1
Tctn1
Kynu
Eml4
Tspan32
Rasgrp1
Arhgef18
Dexi
Lmo2
Xkrx
Ift140
Abcg3
Birc3
Flna
Rps6ka5
Prkacb
Cnn3
Pag1
Cmtm7
Rgl1
Map4k2
Fchsd2
Cyb561d1
Nod1
Rgs18
Dennd1c
Fmnl3
Plcb2
Mum1
Serpinb1c
Noxred1
Sike1
Scai
Nsmce1
Tmem229b
Dennd4b
Prkab2
Rltpr

ESM Figure 5.

CD19$^+$CD138$^-$

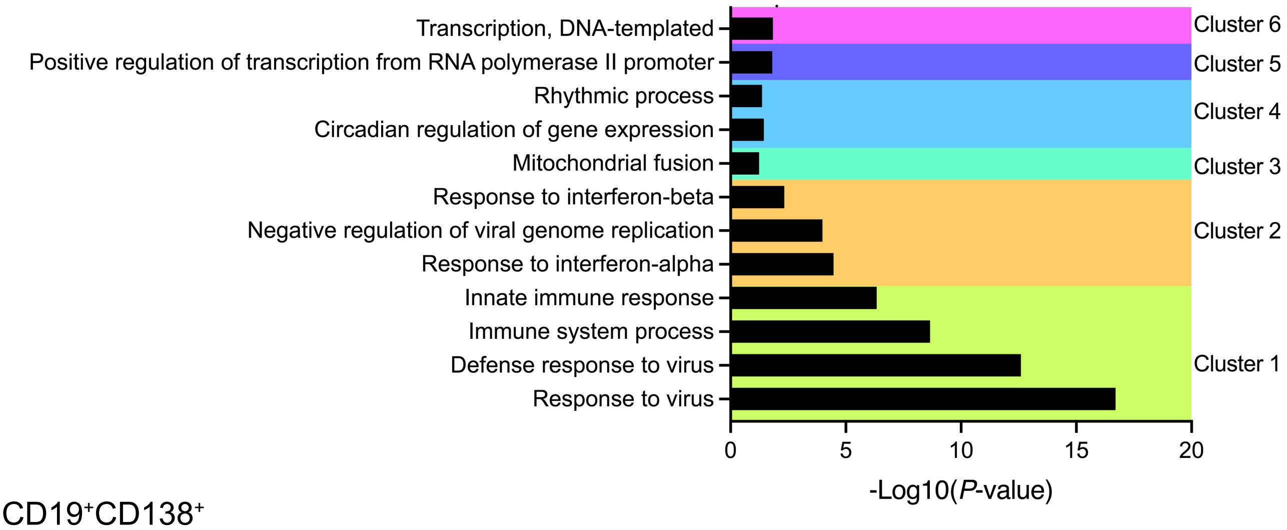


CD19$^+$CD138$^+$

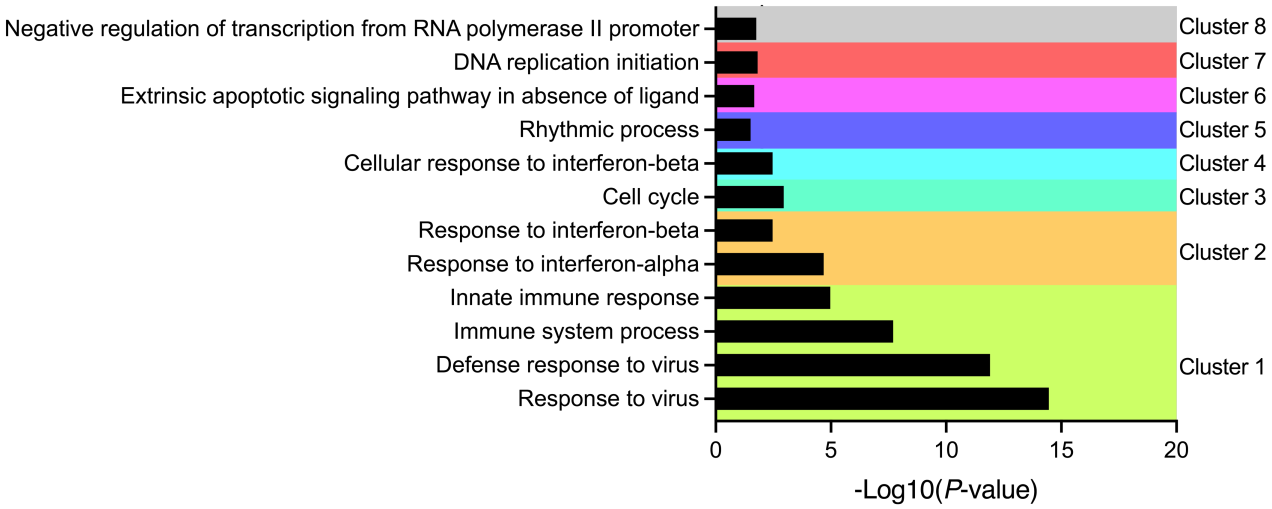

ESM Figure 6.

| |
|---|
| response to virus (GO:0009615) |
| response to external biotic stimulus (GO:0043207) |
| defense response to symbiont (GO:0140546) |
| response to other organism (GO:0051707) |
| defense response to virus (GO:0051607) |
| response to stress (GO:0006950) |
| defense response to other organism (GO:0098542) |
| response to biotic stimulus (GO:0009607) |
| biological process involved in interspecies interaction between organisms (GO:0044419) |
| response to external stimulus (GO:0009605) |
| defense response (GO:0006952) |
| innate immune response (GO:0045087) |
| immune system process (GO:0002376) |
| immune response (GO:0006955) |
| response to interferon-beta (GO:0035456) |
| response to cytokine (GO:0034097) |
| cellular metabolic process (GO:0044237) |
| organic substance biosynthetic process (GO:1901576) |
| response to interferon-alpha (GO:0035455) |
| negative regulation of biological process (GO:0048519) |
| regulation of cell death (GO:0010941) |
| regulation of apoptotic process (GO:0042981) |
| regulation of programmed cell death (GO:0043067) |
| cellular nitrogen compound metabolic process (GO:0034641) |
